# A unique Thy-1-negative immunofibroblast population emerges as a key determinant of fibrotic outcomes to biomaterials

**DOI:** 10.1101/2022.01.09.475583

**Authors:** Daniel Abebayehu, Blaise N. Pfaff, Grace C. Bingham, Surabhi Ghatti, Andrew Miller, Donald R. Griffin, Thomas H. Barker

**Affiliations:** Department of Biomedical Engineering, Schools of Engineering and Medicine, University of Virginia, Charlottesville, VA 22908; Robert Berne Cardiovascular Research Center, School of Medicine, University of Virginia, Charlottesville, VA 22908; Department of Cell Biology, School of Medicine, University of Virginia, Charlottesville, VA 22908

**Keywords:** Biomaterials, fibrosis, fibroblasts, inflammation

## Abstract

Microporous annealed particle (MAP) hydrogels are an exciting new development in biomaterial design. They regulate innate and acquired immunity which has been linked to their ability to evade normal host-material fibrosis. Yet, resident stromal fibroblasts, not immune cells, are the arbiters of the extracellular matrix assembly that characterizes fibrosis. In other idiopathic fibrotic disorders, a fibroblast subpopulation defined by its loss of cell surface Thy-1 expression is strongly correlated with degree of fibrosis. We have previously shown that Thy-1 is a critical αvβ3 integrin regulator that enables normal fibroblast mechanosensing and here, leveraging non-fibrosing MAP gels, we demonstrate that Thy-1^-/-^ mice mount a robust response to MAP gels that remarkably resembles a classical foreign body response. We further find that within the näive, Thy-1^+^ fibroblast population exists a distinct and cryptic αSMA+ Thy-1^-^ population that emerges in response to IL-1β and TNFα. Employing single-cell RNA sequencing, we find that IL-1β/TNFα-induced Thy-1^-^ fibroblasts actually consist of two distinct subpopulations, both of which are strongly pro-inflammatory. These findings illustrate the emergence of a unique pro-inflammatory, pro-fibrotic fibroblast subpopulation that is central to material-associated fibrosis likely through amplifying local inflammatory signaling.

**Significance Statement:** Despite decades of research, implanted biomaterials are still significantly hampered by the foreign body response and fibrotic encapsulation.

Advancements in material design and immunomodulation have made positive impacts, yet, a fuller mechanistic understanding of aberrant ECM remodeling in the biomaterial microenvironment could improve approaches in biomaterial-host interactions. Here, we leverage anti-fibrotic MAP hydrogels and demonstrate that their ability to evade fibrosis is linked to fibroblast Thy-1 (CD90) surface expression. Thy-1^-/-^ mice exhibit elevated NFκB signaling and elevated fibrosis in response to MAP gel implantation. Interestingly, pro-inflammatory cytokines elicit a Thy-1^-^/αSMA^+^ myofibroblast subpopulation. Single-cell RNA-Seq more fully identifies an ‘immunofibroblast’ subpopulation defined by Thy-1 loss and a pro-inflammatory/fibrotic cytokine, chemokine, and cytokine receptor expression profile, suggesting a self-perpetuating pro-fibrotic subpopulation.

## Introduction

Fibrotic encapsulation of implants and the foreign body response lead to biomaterial rejection and most often follow chronic inflammatory responses ^1–4^. Recently, a microporous annealed particle (MAP) hydrogel system was developed that reduces inflammation, evades fibrosis, and accelerates wound closure and regeneration ^5–8^. Given the reduced inflammation and fibrosis associated with MAP gel implantation, understanding how immuno-stromal crosstalk can inspire or mitigate fibrosis could help usher better biomaterial design^9^. In other human fibrotic disorders, Thy-1 negative fibroblasts emerge as a key player in progressive tissue remodeling and fibrosis. This subset of fibroblasts are mechanically ‘agnostic’ and prone to myofibroblastic differentiation in fibrosis in the lung and heart^10–12^. Thy-1 is a GPI-anchored membrane protein that is expressed on fibroblasts, neurons, and thymocytes and we have previously established it is essential to normal fibroblast mechanosensitivity. Thy-1 directly binds αvβ3 integrin *in cis* and both regulates its tonic/baseline activity and primes the integrin for efficient signal transduction. This critical regulatory mechanism for Thy-1 prevents aberrant fibroblast contractility, strain stiffening of surrounding matrix, and fibrosis^13,14^ (Fig 1A).

**Figure 1.**
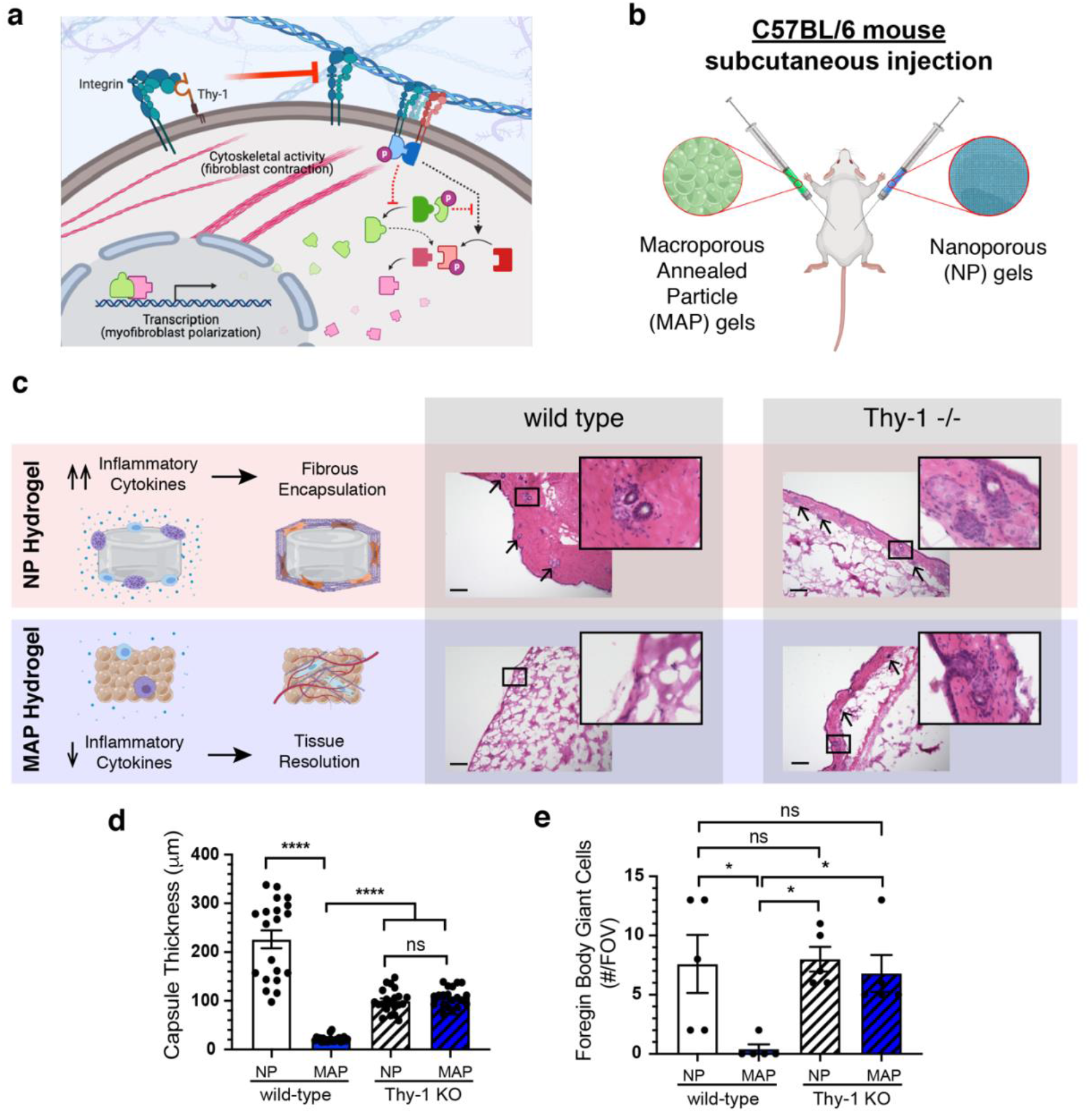
Hydrogel fabrication and subcutaneous implantation. A) Schematic describing regulatory mechanism of Thy-1 in fibroblast mechanotransduction. B) Illustration detailing subcutaneous injection regimen on the dorsal side of mice. C) A summary of tissue and cellular level responses to MAP and nanoporous PEG hydrogels. H&E micrographs of MAP and nanoporous hydrogels subcutaneously implanted into WT and Thy-1^-/-^ mice for 3 weeks, along with inset images to the right providing closer look at fibrotic capsules and the foreign body giant cells present there. D) Quantification of fibrotic capsule thickness and number of foreign body giant cells per field of view in H&E micrographs. **** p<0.0001, * p<0.05, determined with one way ANOVA with Tukey’s multiple comparison test. n = 5 mice per group. Scale bar is 100 microns.

## Results

In order to determine whether fibroblastic expression of the αvβ3 integrin regulator Thy-1 contributes to biomaterial-mediated fibrosis, we specifically asked whether Thy-1 loss is sufficient to reverse the non-fibrotic nature of the host response to MAP hydrogels^5^. We leveraged Thy-1^-/-^ mice and controlled the experiment with a standard fibrogenic nanoporous (NP) bulk hydrogel with the same polyethylene glycol (PEG) chemistry as the implanted MAP gels (Fig 1B). As expected, when implanted subcutaneously into wild-type (WT) mice, the NP hydrogel promoted a thick fibrous capsule after 3 weeks, whereas the MAP hydrogel displayed an almost undetectable fibrous capsule and retained its porous structure, as previously described. In stark contrast, when implanted into Thy-1^-/-^ mice, both hydrogels mounted a robust fibrous encapsulation with no discernable or statistical difference between NP and MAP hydrogels (Fig 1C-D). This key finding is in keeping with prior evidence that fibroblastic loss of Thy-1 expression leads to a highly activated, ECM-synthetic fibroblast, even in normally quiescence-inducing soft environments. Thy-1 negative fibroblasts display high baseline levels of active αvβ3, are far more adhesive and display rapid turnover of integrin adhesion clusters which promotes elevated matrix assembly and contraction, both hallmarks of tissue fibrosis ^12–14^.

Analysis of macrophage-derived foreign body giant cells highlighted a more complex scenario. Specifically, Thy-1 loss was not predicted to impact inflammatory macrophage responses. Again, as expected in WT mice, NP hydrogels promoted a significant number of material-associated foreign body giant cells whereas almost none were associated with MAP hydrogels. However, both NP and MAP hydrogels elicited the same number of foreign body giant cells when implanted in Thy-1^-/-^ mice; a number equivalent to that associated with NP hydrogels implanted into WT mice (Fig 1E). This finding was surprising. Although there is strong evidence to suggest Thy-1^-/-^ fibroblasts are prone to myofibroblastic differentiation and generation of fibrosis, the concomitant elevation in a classical material-associated monocytic-derived immune cells due to a fibroblast defect was unexpected. Furthermore, this immune response resembled a classical reaction to fibrotic materials, implying Thy-1 loss attenuates the beneficial anti-fibrotic properties of MAP gels. These results highlight the significant gap in our understanding of the nuanced relationship between inflammation and fibrosis. Certainly, for over a decade it has been appreciated that skewing macrophage responses can lead to better tissue remodeling outcomes and recently it has been shown that recruitment of IL-33 type 2 myeloid cells and activation of adaptive immunity can drive biomaterial-associated regeneration ^8^. However, these new results suggest that immune signaling may not be unidirectional, from immune cells to stromal cells, but rather bidirectional.

We began unraveling the immuno-stromal signaling axis by exploring IL-1R signaling. Chronic IL-1R signaling is elevated in fibrosis across various tissues ^15–17^. IL-1 receptor I (IL-1RI) expression was clearly evident within the fibrotic capsule of the NP hydrogel and grossly absent in and around the MAP hydrogel within WT mice. In juxtaposition, IL-1RI expression was present and prominent in the fibrotic capsule of both NP and MAP hydrogels implanted in Thy-1^-/-^ mice (Fig 2A). IL-1RI expression within fibrotic capsules strongly correlated with downstream NFκB signaling, as shown by Phosphop-65, and α-smooth muscle actin (α-SMA) expression, a marker of myofibroblast differentiation. Specifically, in NP hydrogels, there was notable co-expression of α-SMA and phosphorylated p65 within the fibrous capsule among WT mice, indicating a potential role for inflammatory signaling in the tissue remodeling phase in addition to the inflammatory phase in response to biomaterials. MAP hydrogels implanted in WT mice demonstrated very little detectable α-SMA expression or p65 phosphorylation at the biomaterial surface or among cells migrating into the implant. Interestingly, these differences are also reflected in regions with less fibrotic encapsulation of the NP hydrogel; there is less αSMA expression and p65 phosphorylation (Supplemental Figure 1A-B). Consistent with our analysis of fibrotic outcomes, NP and MAP hydrogels both display robust α-SMA expression and p65 phosphorylation within their fibrous capsule when implanted in Thy-1^-/-^ mice (Fig 2B-D). The data clearly demonstrate that genetic knock-out of Thy-1 is sufficient to overcome the regenerative and non-fibrotic nature of MAP hydrogels and illustrate a potential fibroblast cytokine signaling mechanism.

**Figure 2.**
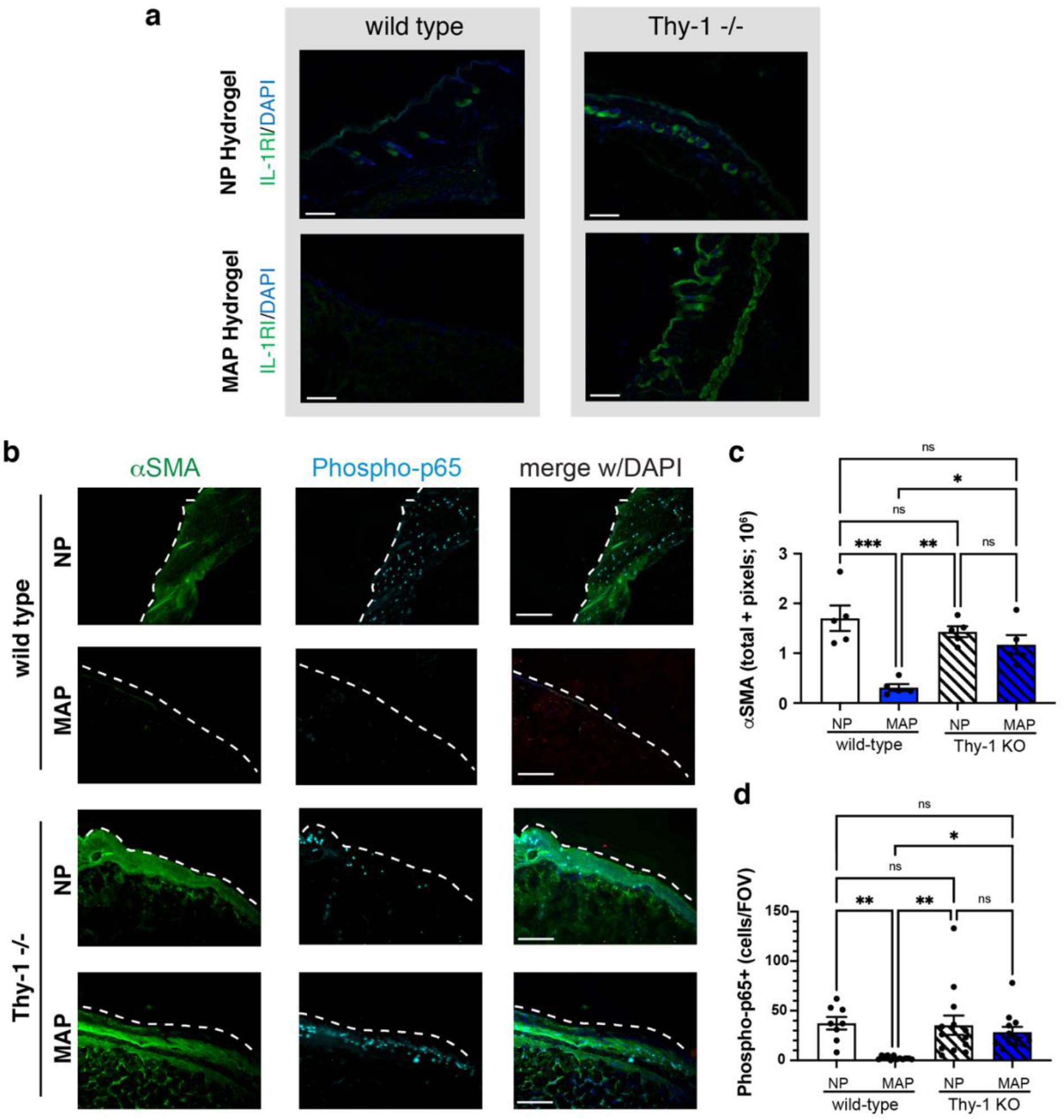
Inflammatory signaling among hydrogels in Thy-1^-/-^ mice. A) Fluorescence images of MAP and nanoporous hydrogel subcutaneous implants stained for IL-1RI (green) and DAPI (blue). B) Fluorescence images staining for αSMA (green), phospho-p65 NFκB (cyan), and DAPI (blue) with quantification for the sum of αSMA pixel values and phospho-p65 NFκB+ cells per field of view. *** p<0.001, ** p<0.01, *p<0.05, determined with one way ANOVA with Tukey’s multiple comparison test. n = 5 mice per group. Scale bar is 100 microns.

The evident differences in immuno-stromal signaling between NP and MAP hydrogels and the ability of Thy-1^-/-^ mice to effectively normalize the fibrotic responses between these two systems led us to hypothesize that inflammatory cytokines promote the loss of Thy-1 and a coincident activated myofibroblastic phenotype among fibroblasts. Using an unbiased approach with a publicly available RNA-Seq dataset (GSM2072332) ^18^ we identified the cytokine receptors expressed on fibroblasts and determined the Th1 (IL-1α, IL-1β, TNFα, INFγ), Th2 (IL-4, −6, −13) and Th17 (IL-17) cytokines that are capable of signaling within fibroblasts. Fibroblasts (CCL-210) treated with Th1 cytokines (IL-1α, IL-1β, INFγ and TNFα) all display elevation in expression of the myofibroblastic marker αSMA (Supplementary Figure 2). We additionally observed significantly elevated fibroblast spreading on soft (2 kPa) fibronectin-coated substrates when treated with IL-1β and TNFα and elevated but not significantly increased spread area when treated with IL-17, INFγ and TGFβ. Only IL-1β and TNFα, and to a lesser extent INFγ, induced a loss of Thy-1 surface expression and only IL-1β and TNFα show consistently promoted αSMA expression. These short-term, acute cytokine stimulations indicate immune-stromal signaling that results in alteration of fibroblast phenotype, but fall short of establishing the emergence of a persistent Thy-1 negative fibroblast subpopulation.

We therefore treated fibroblasts, seeded on soft (~2 kPa) fibronectin coated hydrogels longitudinally with IL-1β and TNFα individually or in combination for 3-7 days. As in short stimulations, IL-1β or TNFα treatment for 72 hours resulted in a significant increase in the Thy-1 negative population (~20% of the total population) compared to fibroblasts in media. The prominence of the Thy-1 negative population was enhanced when stimulated with a cocktail of IL-1β and TNFα (Fig 3A). Stimulation over a 7-day period further intensified the proportion of Thy-1 negative fibroblasts, reaching ~30% in IL-1β and TNFα treatments and the combination yielding fibroblasts that were 40% Thy-1 negative (Supplementary Figure 3). Concomitant with Thy-1 loss, fibroblasts displayed significantly greater cell spreading on soft hydrogels stimulated with IL-1β and TNFα for 72 hours despite showing no discernable differences in focal adhesion size (Fig 3B-C). Fibroblast spreading on soft hydrogels suggests altered mechanosensory signaling and our observations here are consistent with our prior discovery of Thy-1 ‘s role in enabling proper mechanotransduction and stiffness sensing in fibroblasts. Interestingly, only IL-1β stimulation lead to a significant increase in αSMA expression (Fig. 3D), suggestive of a transition to a myofibroblast phenotype. This suggests cooperative, but not overlapping, roles between IL-1β and TNFα signaling in fibroblasts.

**Figure 3.**
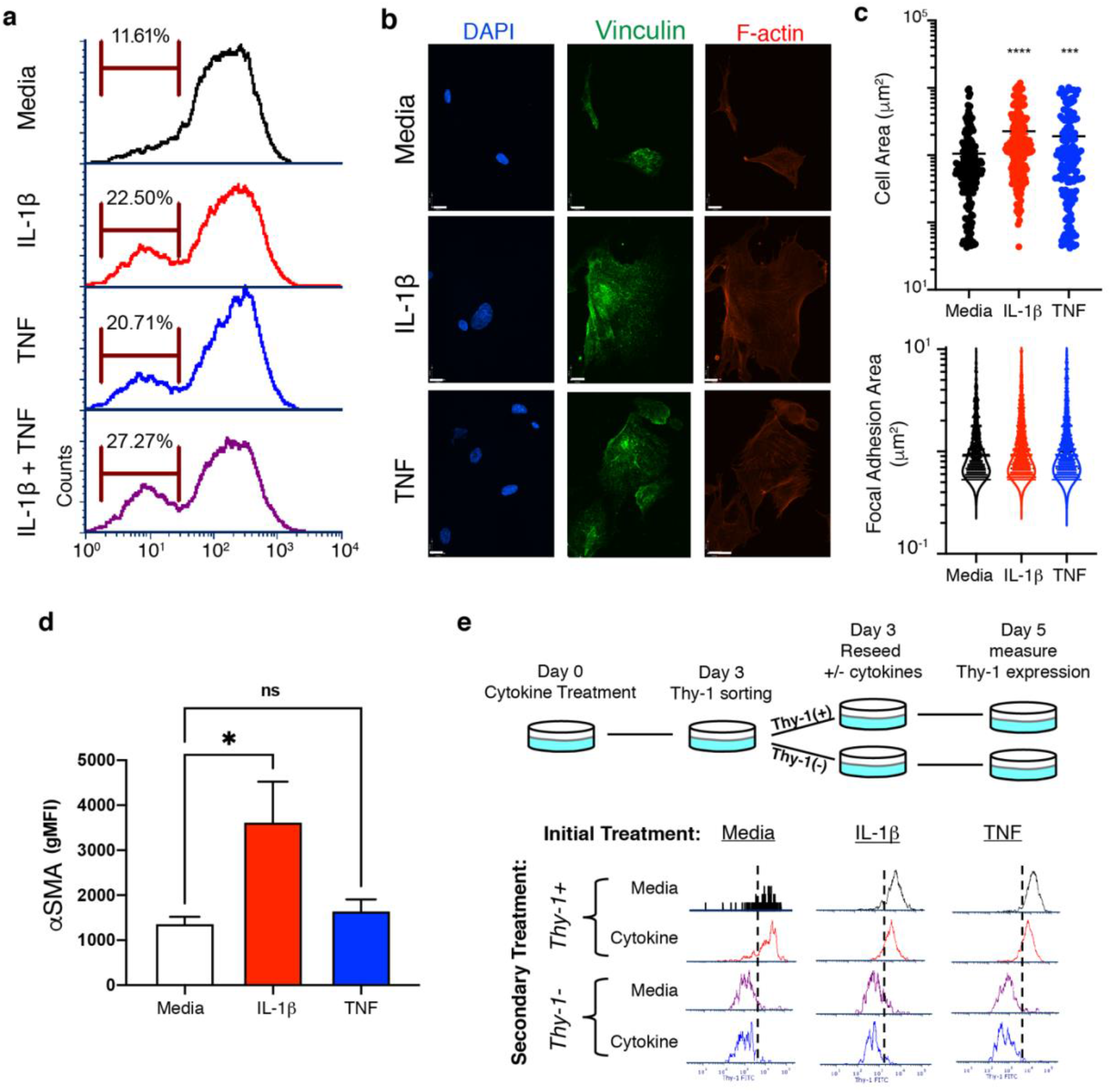
Cytokine-mediated Thy-1 loss. A) Flow cytometric analysis of cytokine treated CCL-210 human lung fibroblasts measuring the emergence of a Thy-1 negative fibroblast subpopulation. B) Fluorescence confocal images of CCL-210 human lung fibroblasts on fibronectin-coated 2 kPa polyacrylamide hydrogels for 48 hours. Staining is for vinculin (green), F-actin (red), and DAPI (blue). C) F-actin images were used to calculate cell area and vinculin images used to measure focal adhesion area. D) Flow cytometric analysis of cytokine treated CCL-210 human lung fibroblasts measuring αSMA expression depicted as gMFI. E) Measuring Thy-1 expression using flow cytometry in CCL-210 human lung fibroblasts that have been sorted based on Thy-1 expression in order to identify whether Thy-1 loss is deterministic. **** p<0.0001, *** p<0.001, * p<0.05, determined with one way ANOVA with Tukey’s multiple comparison test. n = 5 mice per group. Scale bar is 100 microns.

The existence of a true subpopulation implies both a deterministic emergence as well as population persistence. Given the grossly Thy-1 + nature of naïve fibroblasts (only 5-10% are inherently Thy-1-), this would imply that is there a hidden or cryptic fibroblast subpopulation predetermined to lose Thy-1 in response to IL-1β and/or TNFα that then persists in the absence of the stimuli and/or is resistant to additional stimulation. To establish the presence of a deterministic subpopulation, we treated fibroblasts for 72 hours with either IL-1β or TNFα. Fibroblasts were then sorted based on Thy-1 expression into negative and positive populations. Each population was then subsequently stimulated with either media alone or a second dose of the initial cytokine for another 48 hours. Thy-1 + fibroblasts emerging from the initial cytokine stimulation were both stable, i.e. remained Thy-1+ in subsequent culture in media alone, and deterministic, i.e. resistant to Thy-1 loss upon a second stimulation. Similarly, Thy-1-fibroblasts resulting from IL-1β or TNFα stimulation were also stable and deterministic (Fig 3E). These data indicate that there are indeed subpopulations within the native fibroblast population that are either resistant to cytokine-induced loss of Thy-1 and others predisposed to lose Thy-1 and remain Thy-1 negative. The emergent Thy-1-fibroblasts represent a cryptic subpopulation that emerge specifically in response to IL-1β and/or TNFα.

While Thy-1 +/- subpopulations help define the heterogeneity of fibroblasts they certainly do not fully encompass the true heterogeneity within the population. Recently, single-cell RNA sequencing (scRNA-Seq) analysis of fibroblasts from various tissues and diseases has elucidated as many as 5-7 transcriptionally distinct fibroblast subpopulations. We similarly took this approach to uncover both the endogenous heterogeneity of our fibroblast population and how those subpopulations change and new subpopulations emerge (such as Thy-1 -) in response to inflammatory signaling. Our scRNA-Seq analysis revealed 5 transcriptionally distinct clusters of cells among untreated fibroblasts cultured on soft (2 kPa) hydrogels, a degree of heterogeneity that matches what has been observed among fibroblasts in mouse and human tissues ^19–22^. Following 72-hour stimulation with IL-1β and TNF, 2 new clusters emerge (clusters 2 and 5, Fig. 4) along with the expansion of an existing cluster (cluster 0; Fig 4A-E). At the transcriptional level, subpopulations 0, 2, and 5 also lacked Thy-1 expression (Fig 4C). Thus, cluster 0 represents an endogenous, native Thy-1 negative population, whereas clusters 2 and 5 are the emergent, cryptic Thy-1 negative subpopulations observed in prior experiments. Importantly, the top markers of clusters 0, 2, and 5 are inflammation related: IL1RI, IL6, IL1β, IL8, IL33, and IL11 (Supplementary Fig 8A and Fig 4B). These transcriptional signatures were further confirmed by Pathway Enrichment Analysis that identified overexpression of transcripts related to PI3K-Akt, TNF, IL-17, and NFκB signaling pathways (Fig 4D). Intriguingly, the top gene identifying cluster 5 is IL1β, the inflammatory cytokine that induces Thy-1 loss and the emergence of the pro-fibrotic fibroblast subpopulations. Together, these data provide a compelling argument that initial signaling by Th1 cytokines associated with the acute inflammatory response to biomaterials induce a pro-fibrotic immunofibroblast capable of driving a positive feedback autocrine signaling loop capable of sustaining fibrotic progression.

**Figure 4.**
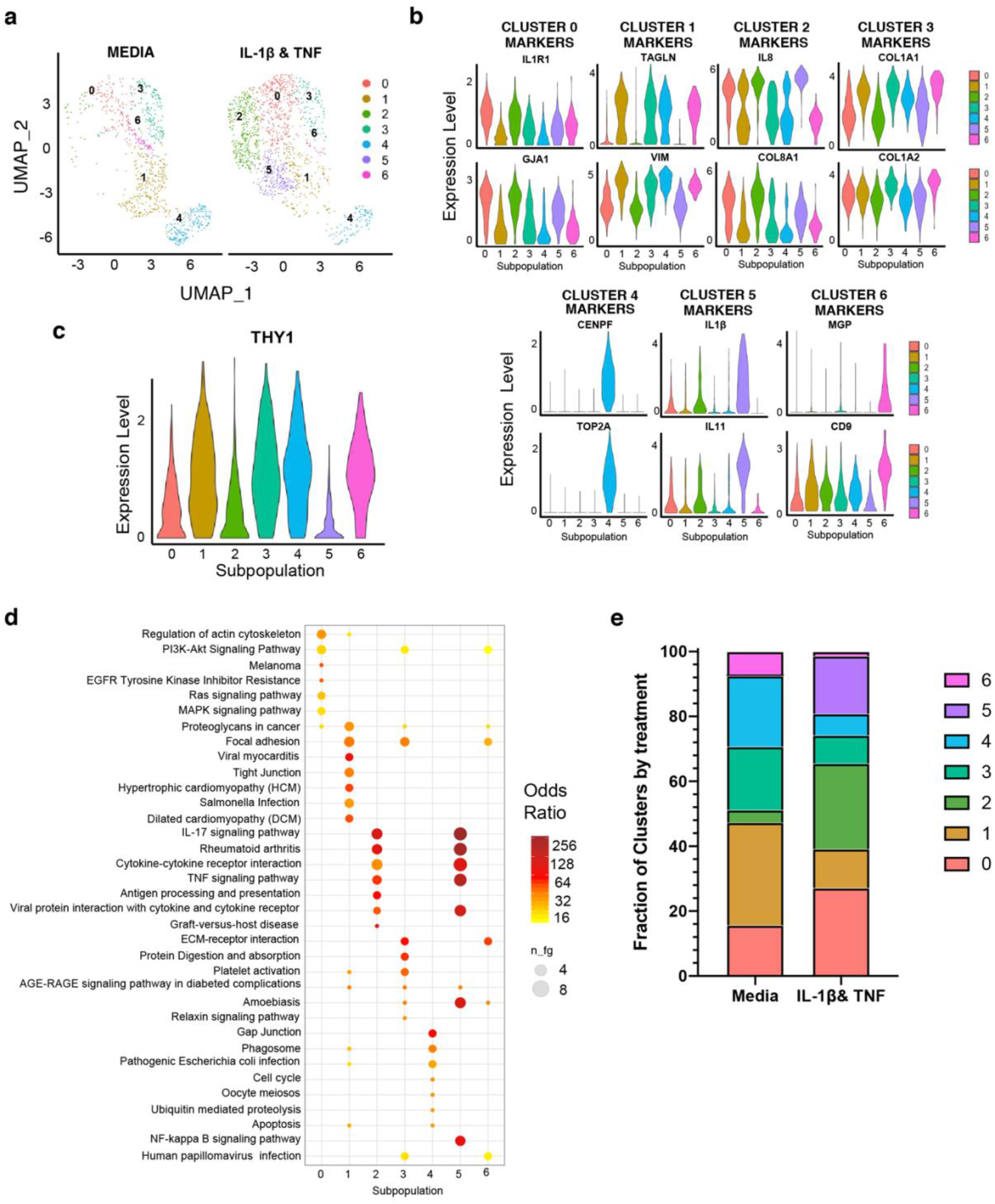
Cytokine-mediated changes in fibroblast heterogeneity. CCL-210s were treated with inflammatory cytokines on 2 kPa fibronectin-coated hydrogels for 72 hours. Cells were collected for scRNA-Seq. A) Uniform Manifold Approximation and Projection (UMAP) clustering projection showing subpopulations of CCL-210 fibroblasts in media alone or treated with IL-1β and TNFα. B) Violin plots of top 2 differentially expressed genes that serve as markers identified for each subpopulation. C) Violin plot demonstrating Thy-1 expression across subpopulations. D) KEGG pathway enrichment analysis identifying upregulated pathways corresponding to differentially expressed genes for each cluster. E) Fraction of cells coming from each subpopulation based on treatment group.

## Discussion

Biomaterial-mediated fibrosis and the foreign body response are the critical obstacles to implant success. While inflammatory signaling and various immune cell populations are critical to fibrotic encapsulation, yet it is the fibroblasts and not immune cells that are the primary contributors to scar formation. Therefore, it is critical to understand how immune-fibroblast crosstalk contributes to biomaterial-mediated fibrosis. Here we demonstrate that implants that undergo fibrotic encapsulation display robust NFκB signaling as well as IL-1RI+ fibroblasts within the fibrous capsule. We demonstrate that IL-1β treatment promoted αSMA expression, increased cell area indicative of increased cell contractility, and loss of Thy-1 expression. Thy-1 loss has been shown to be critical to various forms of tissue fibrosis and promotes a mechanically agnostic phenotype among fibroblast subpopulations that undergo myofibroblastic differentiation independent of the mechanical cues from the surrounding microenvironment. Interestingly, when non-fibrotic and regenerative MAP gels were implanted in Thy-1^-/-^ mice, these MAP gels induced a fibrotic response that resembled the encapsulation seen in response to bulk nonporous gels. Our data show that the ability to evade fibrotic remodeling by MAP gels is not necessarily a salient feature of the material, per se, and can be reversed by simply tuning the fibroblast cell milieu within the material microenvironment. scRNA-Seq determined that IL-1β treatment altered the fibroblast subpopulations and promoted the emergence of a Thy-1 negative ‘immunofibroblast’ defined by cytokine, chemokine, and cytokine receptor expression.

Macrophage/monocyte-to-myofibroblast transitions ^23^, so-called ‘fibrocytes’, have been well described in fibrotic responses. Here, in a surprising twist, elements of the transcriptional signature of these unique fibroblasts include traditional macrophage markers (Supplementary Figure 8B), implicating a form of myofibroblast-to-macrophage transition such that fibroblasts are capable of sustaining a macrophage-like response well beyond the termination of the acute inflammatory phase. Immuno-stromal communication has been previously reported in disease ^24^ and endogenous repair ^25^ where inflammation instructs fibrosis, yet our data suggest that fibroblasts can “talk back” driving chronic inflammation. Tackling these unexpected and cryptic contributors to inflammation and fibrosis will be the key to future success in biomaterial development and the longevity of implant function.

## Methods

### Hydrogel Fabrication

A 6.5wt% gel with a PEG-Maleimide backbone was prepared by dissolving PEG-Mal, MethMal, and RGD in pH=2 10X PBS. This solution was combined with an MMP crosslinker and 5μM biotin-maleimide which were dissolved in ultrapure water. The final concentrations in the hydrogel solution were 81.31mg/mL PEG-Mal, 14.90mg/mL MMP, 0.80mg/mL RGD, and 8.02mg/mL of MethMal. Microgels were produced using a microfluidics PDMS mold created using previously published design^26^ in a dust free hood. Briefly, a 1% Pico-Surf surfactant (Sphere Fluidics) solution diluted in NOVEC 7500 oil (3M) was run through the oil channel and the gel formulations described above were in the aqueous channel. Using a syringe pump, the surfactant and gel solutions were run at 5mL/hr in the device and collected in a conical tube. The resultant particles were mixed with a triethylamine solution (20μL TEA/ mL of gel) to increase the pH and accelerate gelation. Microgels were washed with NOVEC 7500 oil three times (1X volume of gel). Next, microgels were combined with PBS (5X volume of gel) and washed with NOVEC 7500 oil (1X volume of gel) three times, allowing separation of the oil and aqueous solution by settling. Finally, the oil was removed and the microgels were washed with PBS (3X volume of gel) and hexanes (3X volume of gel) three times. After the final wash, microgels reacted overnight at 37°C with a 100mM N-Acetyl-L-Cysteine solution in 1X PBS to cap any excess maleimides. Microgels were removed from solution via centrifugation (4696 gx5min) and all further steps were performed in a biosafety cabinet. Microgels were washed three times with 70% isopropyl alcohol followed by four washes with sterile 1X PBS (pH=7.4) before being mixed with a photoinitiator solution.

### Nanoporous Gel Fabrication

In a biosafety cabinet, an identical 6.5wt% gel with a PEG-Maleimide backbone was prepared by dissolving PEG-Mal, MethMal, and RGD in sterile-filtered pH=6.74 2X PBS. The MMP crosslinker and 5μM biotin-maleimide were dissolved in ultrapure water. The solutions were combined and loaded into a syringe for injection. The final concentrations in the nanoporous gel solution were 81.31 mg/mL PEG-Mal, 14.90mg/mL MMP, 0.80mg/mL RGD, and 8.02mg/mL of MethMal.

#### Subcutaneous Implantation of Hydrogels

For subcutaneous implantation of hydrogels, 10-week-old C57BL/6 mice were anesthetized with aerosolized isoflurane and received three 80uL subcutaneous dorsal-side injections of MAP hydrogel solution and three 80uL injections of NP hydrogel solution as a contralateral control. After 3 weeks, mice were euthanized via carbon dioxide asphyxiation and the hydrogels and surrounding subcutaneous tissue were collected and frozen in OCT over dry ice. 10um sections were cut using a CryoStar NX50 cryostat and mounted on SuperFrost Plus slides for downstream histology and immunofluorescence.

#### Histology and Analysis

Slide-mounted 10um sections of hydrogel samples were fixed with 4% paraformaldehyde and then stained with H&E using standard procedures. The subsequent histology images were blinded and analyzed for fibrous capsule thickness, which we defined as the collagenous tissue surrounding the circumference of the hydrogel implant, and foreign body giant cell presence, which we defined as multinucleated cells within the fibrotic capsule with a large bundle or ring of nuclei ^27^.

#### Tissue Immunofluorescence and Quantification

Slide-mounted 10um sections were fixed with 4% paraformaldehyde and blocked with 3% normal goat serum (NGS) + 0.1% Triton in 1XPBS for intracellular antigens. Primary antibody dilutions were prepared in 3% NGS in 1XPBS as follows: anti-mouse α-SMA (1A4, ThermoFisher) at 1:200, rat anti-mouse Thy-1.2 (53-2.1, BD Biosciences) at 1:100, rat anti-mouse phospho-NFκB p65 (Ser536; 93H1, Cell Signaling Technologies) at 1:100, and biotinylated anti-mouse IL-1RI (JAMA-147, BioLegend) at 1:100. The primary antibody incubation was done overnight at 4°C, then three 1XPBS washes, and then a 1-hour secondary antibody staining at room temperature. The secondary antibodies (goat anti-mouse Alexa Fluor 488, goat anti-rat Alexa Fluor 555, goat anti-rabbit Alexa Fluor 647; Invitrogen) were all prepared in 3% NGS in 1XPBS at a dilution of 1:1000. DAPI was used as a counterstain at 300nM in 1xPBS for 5 minutes at room temperature and then mounted in mounting medium ProLong Diamond Antifade (ThermoFisher). Slides were imaged on a Keyence BZ-X810 microscope using the 10X objective. α-SMA expression was quantified by defining a region of interest in Fiji around the fibrotic capsule within the field of view and measuring the sum of the α-SMA signal pixel intensity as defined by Raw Integrated Density in Fiji. Phospho-p65+ cells were measured by recording the number of cells that expressed phosphop-p65 in thresholded images per field of view using the “Analyze Particles” function in Fiji.

#### Polyacrylamide Hydrogels for Cell Culture

Human lung fibroblasts (CCL-210) were purchased from ATCC were cultured (DMEM, 10% fetal bovine serum, 1% penicillin/streptomycin) and used between passage 2-10. To recapitulate the biophysical microenvironment and remove substrate stiffness as a conflating factor in this study, fibroblasts were cultured on 2 kPa polyacrylamide hydrogels that were purchased from Matrigen and came chemically activated ready to bind matrix proteins, and were subsequently coated in 10ug/mL of fibronectin. For immunocytochemistry of fibroblasts on hydrogels, cells were fixed with 4% paraformaldehyde and blocked with 3% normal goat serum (NGS) + 0.1% Triton in 1XPBS for intracellular antigens. Primary antibody dilutions were prepared in 3% NGS in 1XPBS as follows: mouse anti-human vinculin (VIN-54, Abcam) at 1:200 and phalloidin conjugated with Alexa Fluor 546 (Invitrogen). The primary antibody incubation was done for 1 hour at room temperature, then three 1XPBS washes, and then a 1-hour secondary antibody staining at room temperature. The secondary antibody (goat anti-mouse Alexa Fluor 488, Invitrogen) was all prepared in 3% NGS in 1XPBS at a dilution of 1:1000. DAPI was used as a counterstain at 300nM in 1xPBS for 5 minutes at room temperature and then mounted in mounting medium ProLong Diamond Antifade (ThermoFisher). Imaging was done at room temperature on a Nikon Eclipse Ti microscope with an UltraView VoX imaging system (PerkinElmer) using a Nikon N Apo LWD 40X water objective (numerical aperature: 1.15). Cell and focal adhesion area were measured using Fiji software.

#### Flow Cytometry and FACS Sorting

Fibroblasts were lifted from hydrogels for flow cytometry using TrypLE Express Enzyme (ThermoFisher) and subsequently stained with the Zombie Near-Infrared Fixable Viability dye (BioLegend) then fixed and permeabilized using the FIX & PERM Cell Permeabilization Kit (ThermoFisher). After fixation, fibroblasts were surface stained for Thy-1 using the APC anti-Thy-1 antibody (1:100, BioLegend). For α-SMA staining, after surface staining and washing, fibroblasts were incubated in the permeabilization buffer with the PE anti-α-SMA antibody (1A4, R&D) at a 1:25 dilution. Cells were then washed and analyzed on a BD LSR Fortessa through the UVA Flow Cytometry Core Facility and analysis done using FCS Express 7.

#### Single Cell RNA-Sequencing and Analysis

Fibroblasts were lifted from hydrogels using TrypLE Express Enzyme and viability was confirmed to be greater than 80% using Trypan blue staining with the Countess II FL Automated Cell Counter (ThermoFisher). Cells were resuspended in 0.04% UltraPure BSA in PBS (ThermoFisher). 1,500 fibroblasts in each group were targeted. After 72 hours of cytokine treatment, cells will be collected and then we used the 10X Genomics Chromium platform for automated single cell barcoding and library preparation on an 8-channel microfluidics chip. Then we used the Illumina NextSeq 500 Sequencing System for high throughput sequencing that allows for 400M reads, permitting us to have broad coverage of each cell’s transcriptome. The Genome Analysis core at UVA assisted in single cell barcoding, library construction, and sequencing. Gene-barcode matrices were analyzed in R using Seurat v3 ^28^. Cells were filtered for 2,500 to 9,000 reads per unique molecular identifiers and less than 10% of mitochondrial gene content. Significant principal component analysis of variation was calculated using JackStraw test with 100 repetitions and clusters were defined using 20 principal components of variation. Code is available on request.

#### Statistical Analysis

Statistics were performed using GraphPad Prism 9 (GraphPad, San Diego, CA). For multiple group comparisons, we performed 1-way ANOVA followed by Tukey’s post hoc test. The number of mice used for each analysis was determined by power analysis (power = 0.80, α = 0.05). Statistical significance was defined at p-values < 0.05.

## Data Availability

Data will be made available upon request.

## Acknowledgements

We acknowledge the University of Virginia Flow Cytometry Core Facility for assistance with sorting and access to flow cytometers. We also want to acknowledge the University of Virginia Genome Analysis and Technology Core for assistance with single cell RNA-Sequencing library prep, sequencing, and alignment of raw data. Figures 1A and 1B are schematics created with BioRender.com. D.A. acknowledges financial support from the U.S. National Heart, Lung, and Blood Institute fellowship (5F32HL147405-02) and from the U.S. National Institutes of Health Cardiovascular Training Grant (2T32HL007284-41). B.N.F. and A.M. acknowledge funding from the U.S. National Institutes of Health Biotechnology Training Grant (1T32GM136615-01). G.C.B. acknowledges funding from the U.S. National Institutes of Health Cancer Training Grant (5T32CA009109-44). D.R.G. acknowledges funding from the U.S. National Institutes of Health High Priority, Short-Term Project Award (1R56DK126020-01). T.H.B. acknowledges funding from the U.S. National Institutes of Health (5R01HL132585-05 and 5R01HL127283-05).

## Author Contributions

D.A. and T.H.B. designed the experiments. B.N.F. performed biomaterial fabrication and implantation. D.A. performed immunofluorescence and histological staining on implanted biomaterials, in vitro experiments measuring Thy-1 loss and its dynamics, measuring cell contractility and focal adhesions, and single cell RNA-Sequencing. G.C.B. performed immunocytochemistry experiments with fibroblasts on hydrogels. S.B. performed in vitro experiments measuring αSMA via flow cytometry. D.A. and A.M. performed analysis on single cell RNA-Seq dataset. D.A. performed all experimental analysis and statistical tests. D.A., B.N.P., G.C.B., S.G., A.M., D.G., and T.H.B. wrote the manuscript.

---

**Supplemental Figure 1.**
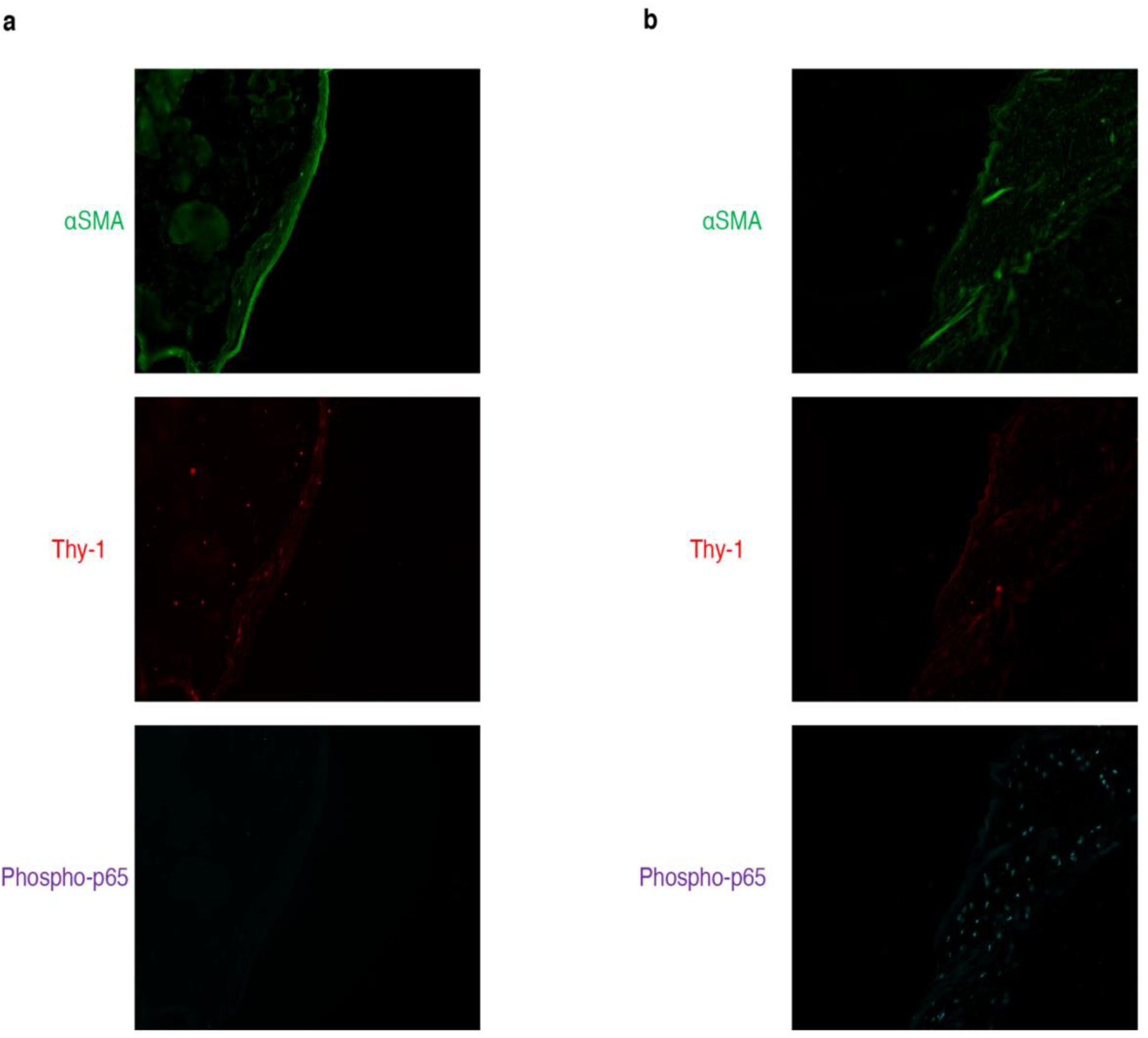
Fibrotic and non-fibrotic regions of nanoporous hydrogels. A) Region of nanoporous hydrogel with minimal fibrous capsule formation and immunofluorescence staining for αSMA, Thy-1, and phospho-p65 NFκB. B) Region of nanoporous hydrogel with robust fibrous capsule formation and immunofluorescence staining for αSMA, Thy-1, and phospho-p65 NFκB.

**Supplemental Figure 2.**
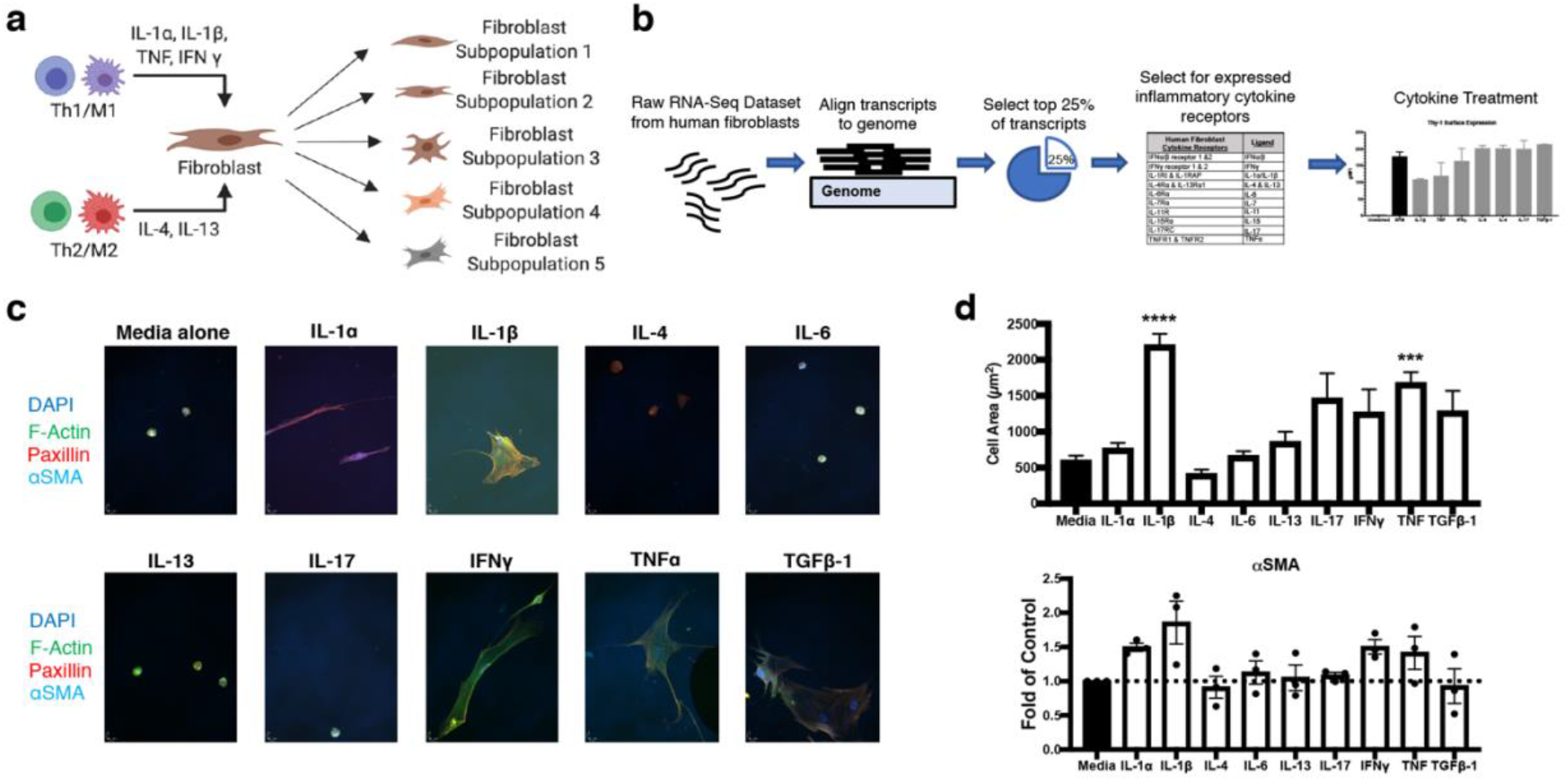
Identifying cytokine receptors and cytokine panel. A) Schematic illustrating how soluble factors from different immune cell subpopulations may promote emergence of fibroblast subsets with varying phenotypes. B) Analysis pipeline of publicly available normal human lung fibroblast RNA-Seq dataset for identifying human lung fibroblast cytokine receptor expression. Table shows the panel of receptors identified and the corresponding ligands included in our cytokine panel. Chart to the right shows CCL-210 Thy-1 loss in response to the panel of cytokines. C) Immunocytochemistry of CCL-210s on fibronectin-coated 2 kPa polyacrylamide hydrogels staining to assess cytoskeletal remodeling and myofibroblastic differentiation in response to cytokine treatment. D) Cell area, measured based on F-actin staining, and αSMA expression were analyzed to determine which of the cytokines in the panel recapitulated the Thy-1 negative phenotype previously reported.

**Supplemental Figure 3.**
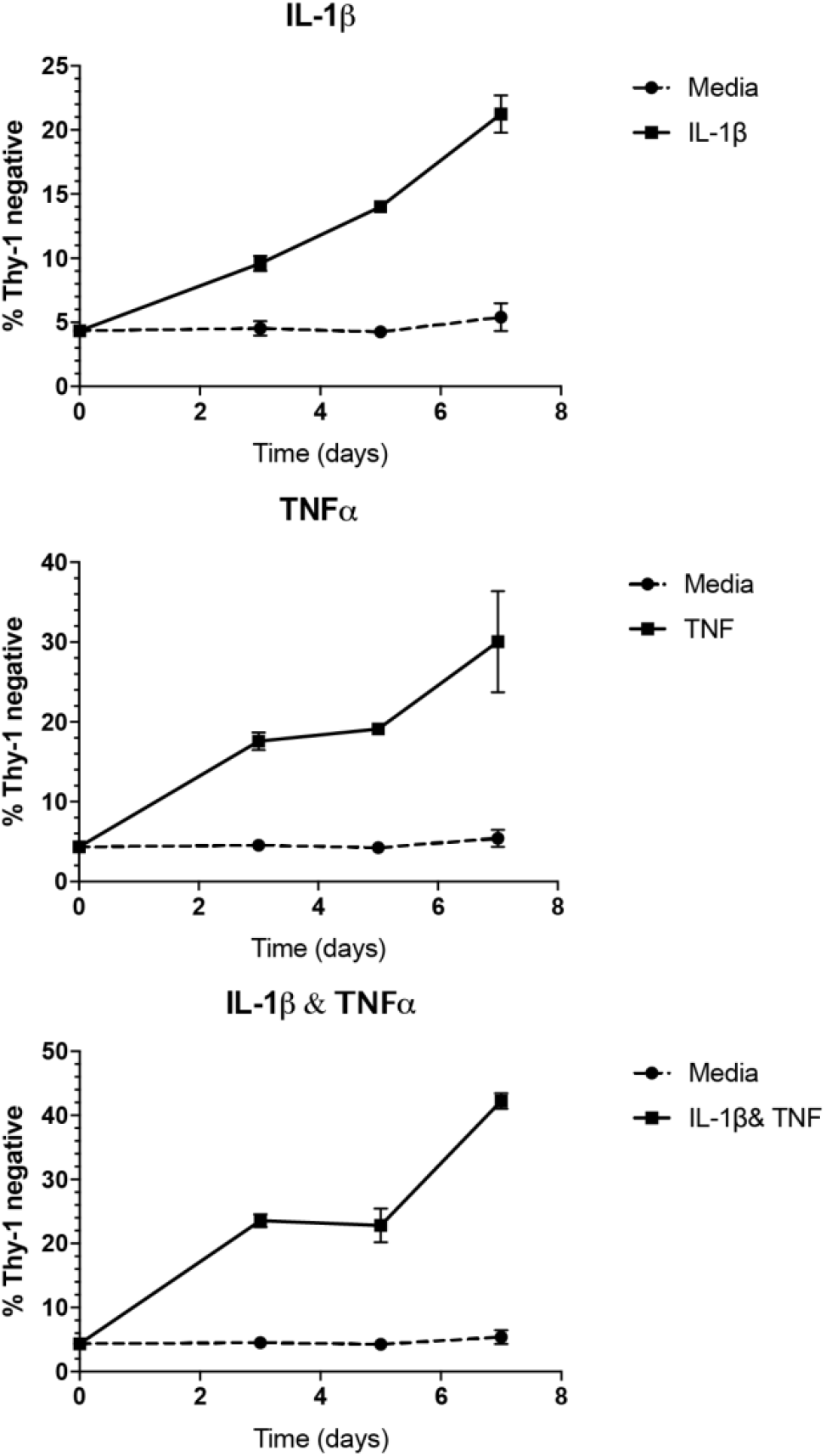
Time-course for Thy-1 loss in response to cytokines. CCL-210s were seeded on fibronectin-coated 2 kPa polyacrylamide hydrogels in the presence of cytokines (all at 50ng/mL) for 3, 5, and 7 days and the emergence of a Thy-1 negative subpopulation was measured.

**Supplemental Figure 4.**
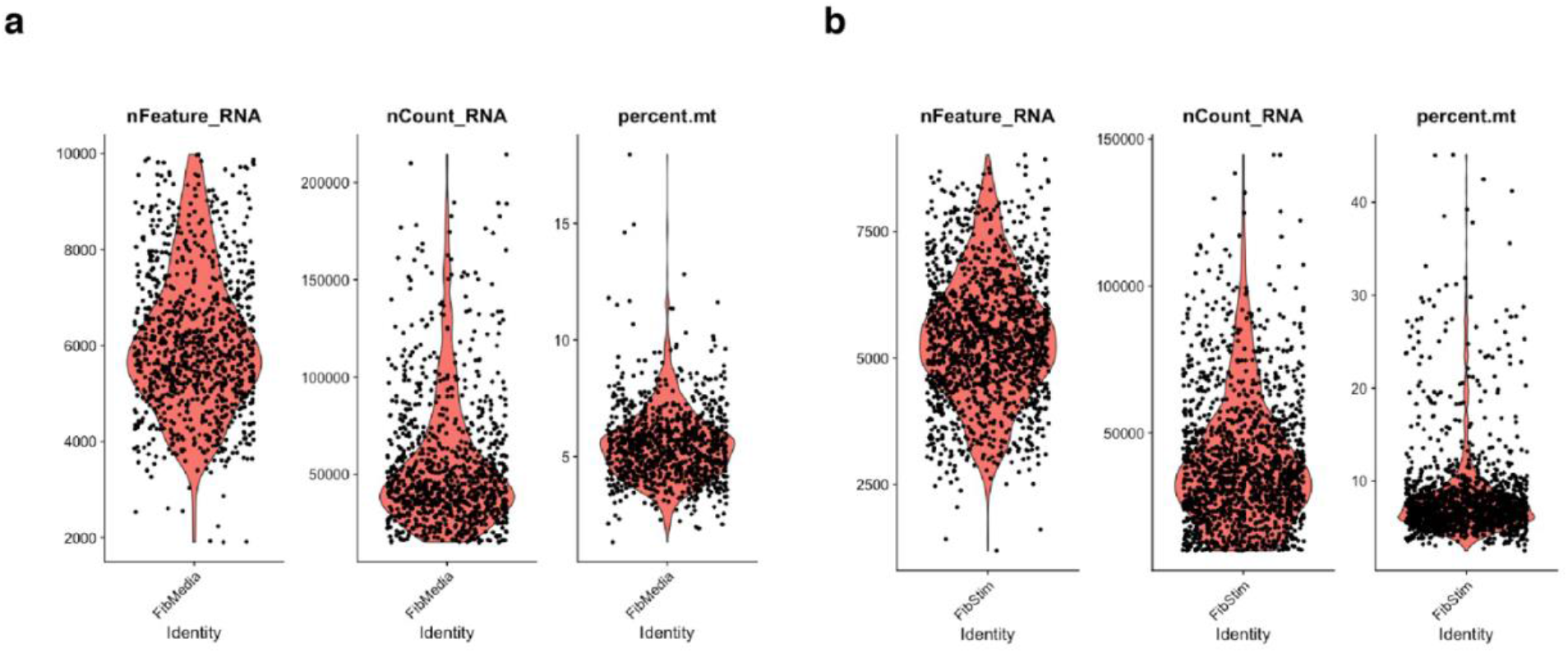
scRNA-Seq QC. These violin plots demonstrate the distribution of cells and how many unique molecular identifiers (UMIs) were calculated per cell as well as the mitochondrial gene content in the A) media alone and B) cytokine treated samples. Based on these plots, cells were filtered for 2,500 to 9,000 reads per unique molecular identifiers and less than 10% of mitochondrial gene content.

**Supplemental Figure 5.**
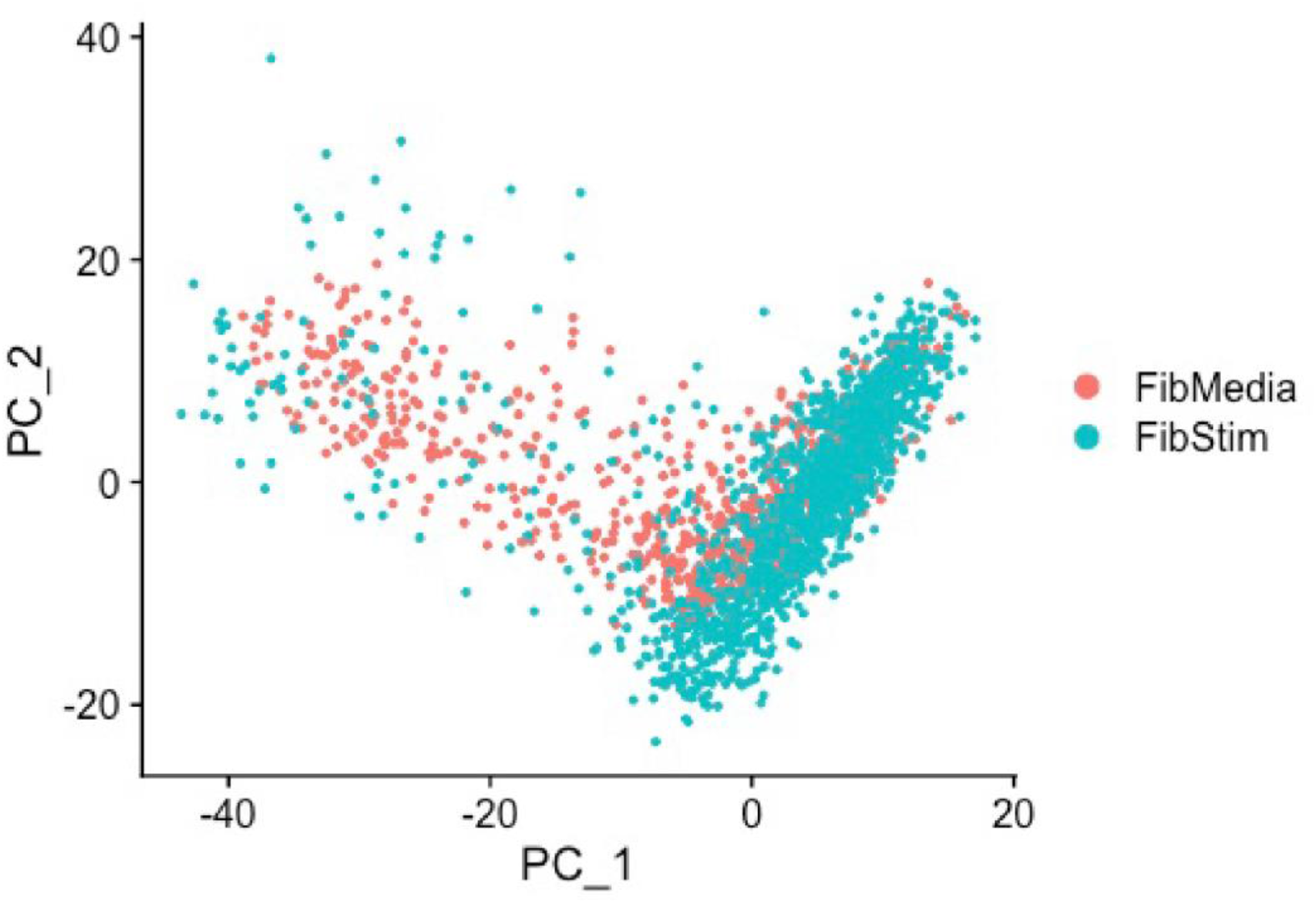
PCA Dimensionality Reduction. After QC, the scRNA-Seq dataset underwent PCA dimensionality reduction to identify the variance between the media and cytokine treated fibroblast samples.

**Supplemental Figure 6.**
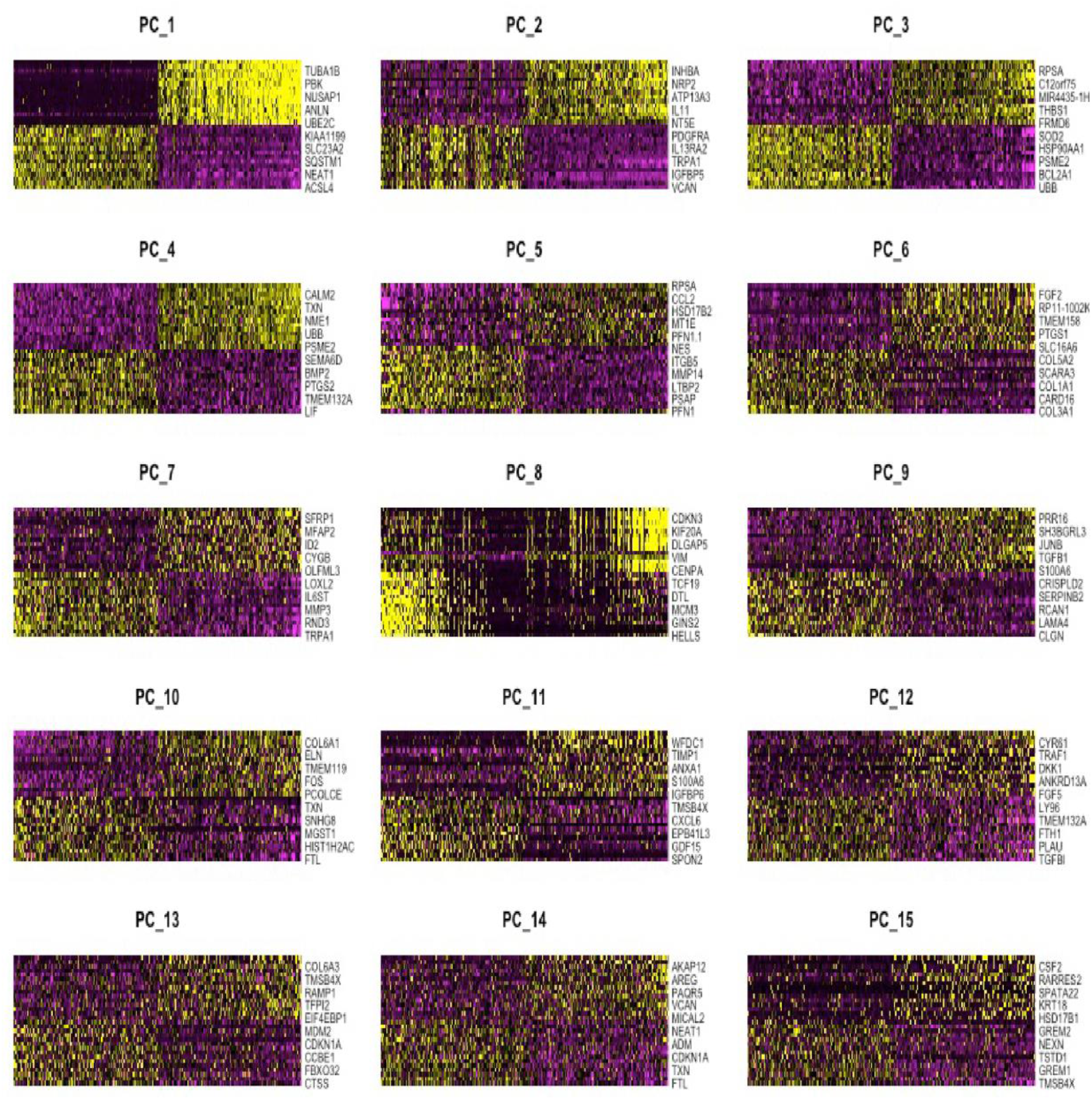
Heatmaps to determine heterogeneity. A series of heatmaps were created for all identified PCA groups for the integrated dataset to determine the unique markers for each group.

**Supplemental Figure 7.**
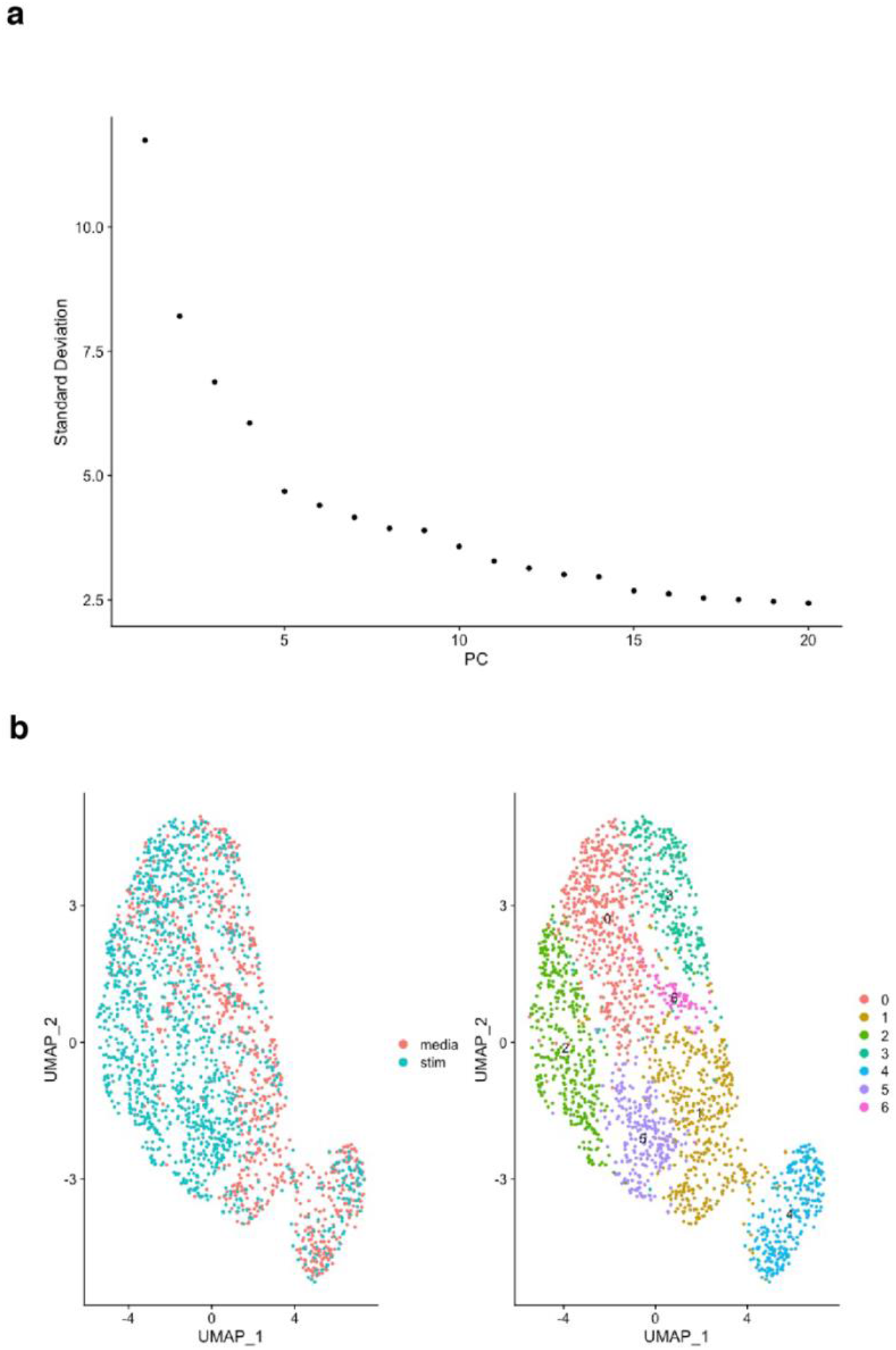
Cluster Identification. A) JackStraw plot was created to identify the optimal number of clusters that best characterizes the heterogeneity within the dataset. B) UMAP clustering projection of the scRNA-Seq dataset separated by treatment groups (media alone and cytokine treatment) as well as a UMAP plot of the 7 clusters of the integrated dataset.

**Supplemental Figure 8.**
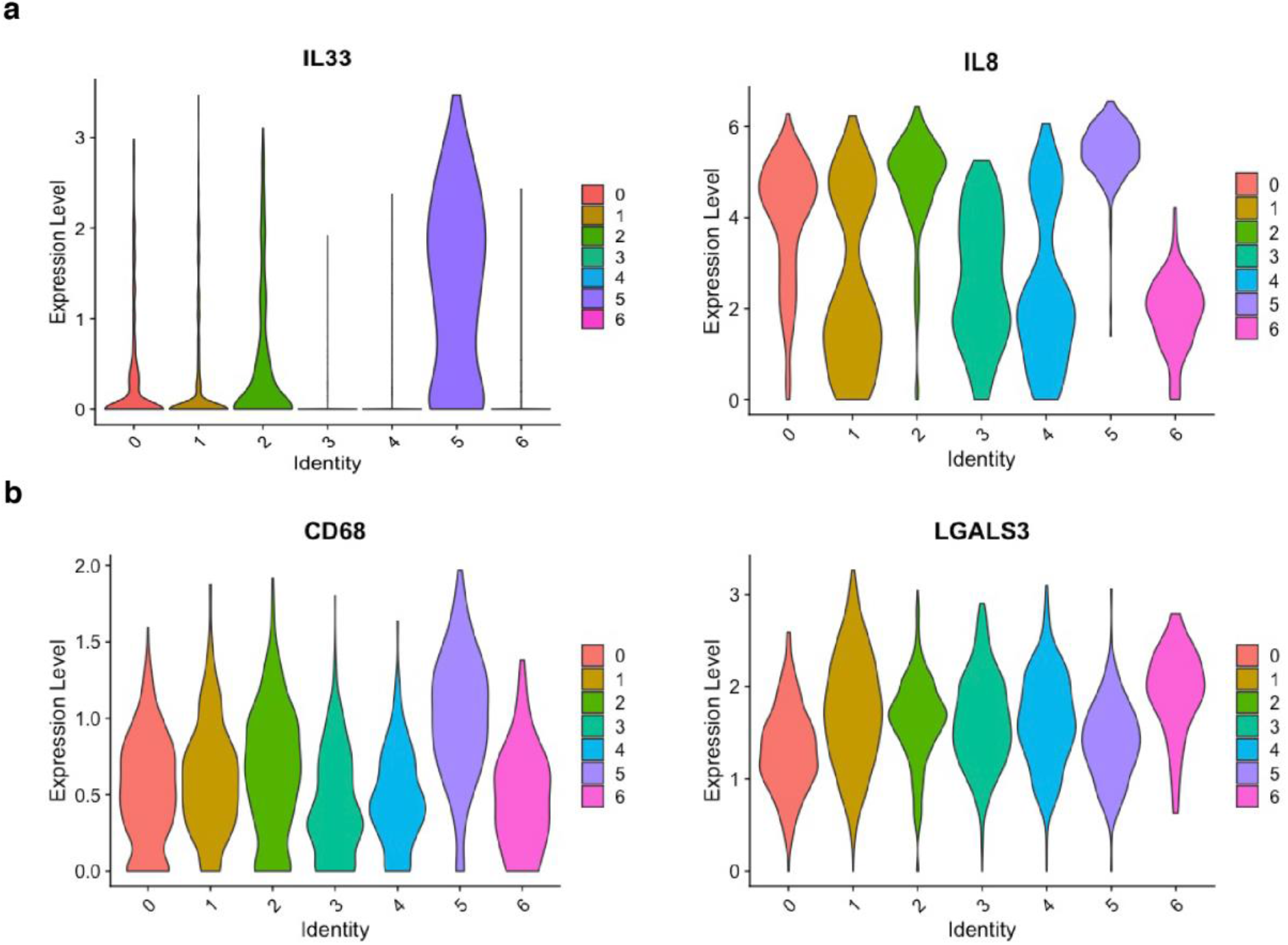
Inflammatory and immune cell markers. A) Violin plots of inflammatory cytokines that are top markers for clusters 2 and 5, the clusters that are also are lowest in Thy-1 transcript expression. B) Interestingly, the top surface markers for clusters 2 and 5 are coincidentally also traditionally considered to be top macrophage markers, indicating a potential macrophage-like phenotype among the fibroblasts in these subpopulations.

